# Complete genomes of clade G6 Saccharibacteria suggest a divergent ecological niche and lifestyle

**DOI:** 10.1101/2021.05.28.446221

**Authors:** Jonathon L. Baker

## Abstract

Saccharibacteria (formerly TM7) have reduced genomes, a small size, and appear to have a parasitic lifestyle dependent on a bacterial host. Although there are at least 6 major clades of Saccharibacteria inhabiting the human oral cavity, cultured isolates or complete genomes of oral Saccharibacteria have been previously limited to the G1 clade. In this study, nanopore sequencing was used to obtain three complete genome sequences from clade G6. Phylogenetic analysis suggested the presence of at least 3-5 distinct species within G6, with two discrete taxa represented by the 3 complete genomes. G6 Saccharibacteria were highly divergent from the more well-studied clade G1, and had the smallest genomes and lowest GC-content of all Saccharibacteria. Pangenome analysis showed that although 97% of shared pan-Saccharibacteria core genes and 89% of G1-specific Core Genes had putative functions, only 50% of the 244 G6-specific Core Genes had putative functions, highlighting the novelty of this group. Compared to G1, G6 encoded divergent metabolic pathways. G6 genomes lacked an F1F0 ATPase, the pentose phosphate pathway, and several genes involved in nucleotide metabolism, which were all core genes for G1. G6 genomes were also unique compared to G1 in that they encoded lactate dehydrogenase, adenylate cyclase, limited glycerolipid metabolism, a homolog to a lipoarabinomannan biosynthesis enzyme, and the means to degrade starch. These differences at key metabolic steps suggest a distinct lifestyle and ecological niche for clade G6, possibly with alternative hosts and/or host-dependencies, which would have significant ecological, evolutionary, and likely pathogenic, implications.

**Importance:** Saccharibacteria are ultrasmall, parasitic bacteria that are common members of the oral microbiota and have been increasingly linked to disease and inflammation. However, the lifestyle and impact on human health of Saccharibacteria remains poorly understood, especially for the 5 clades (G2-G6) with no complete genomes or cultured isolates. Obtaining complete genomes is of particular importance for Saccharibacteria, because they lack many of the “essential” core genes used for determining draft genome completeness and few references exist outside of clade G1. In this study, complete genomes of 3 G6 strains, representing two candidate species, were obtained and analyzed. The G6 genomes were highly divergent from G1, and enigmatic, with 50% of the G6 core genes having no putative functions. The significant difference in encoded functional pathways is suggestive of a distinct lifestyle and ecological niche, probably with alternative hosts and/or host-dependencies, which would have major implications in ecology, evolution, and pathogenesis.

## Observation

Saccharibacteria (formerly TM7) have an ultrasmall cell size, reduced genomes, and are thought to be obligate epibionts, dependent on physically-associated host species (1-3). Common constituents of the oral microbiota, Saccharibacteria have been increasingly linked to inflammation and disease (4-6). Saccharibacteria contains at least 6 distinct clades (G1-G6)(7, 8), however all currently available human-associated complete genomes and cultured isolates belong to clade G1, leaving clades G2-G6 quite poorly understood. Several recent publications have provided the first draft genomes from clades G3, G5, and G6 (4, 8-11). Obtaining complete genomes is of particular importance for Saccharibacteria, because they lack many of the “essential” single-copy core genes that are typically used to estimate genome completion, as well as complete reference genomes outside of the G1 clade.

A recent, short-read-based oral microbiome study provided 21 Saccharibacteria draft genomes from clades G1, G3, and G6 (4), with several being high quality (high N50, relatively contiguous, low predicted contamination). Therefore, nanopore sequencing of the same saliva samples that had produced the draft genomes, followed by long-read and/or hybrid assembly, was used to improve these genomes, resulting in 3 complete, circular G6 genomes: JB001 (662,051 bp), JB002 (639,751 bp), and JB003 (663,165 bp). Table 1 is a summary of the genomes improved during this study and the Supplemental Methods contain a full description of the DNA extraction, sequencing, assembly, and analysis methods. These methods are a modified version of a previously reported protocol (Baker 2021, in-press). Although the G1 and G3 “near complete” improved genomes that were obtained are useful in their own right, they are still incomplete, and/or may contain contamination, therefore the 3 complete G6 genomes are the focus of this report, and the near complete genomes are briefly discussed in the Supplemental Methods.

**Table 1.** Saccharibacteria genomes improved using nanopore sequencing in this study.

| <u>New MAG designation</u> | <u>Previous MAG name</u> | <u>previous # of contigs</u> | <u>previous size (bp)</u> | <u>updated # of contigs</u> | <u>updated size (bp)</u> | <u>updated longest contig (bp)</u> | <u>complete</u> | <u>near complete (longest contig > 700,000 bp or < 5 contigs)</u> |
| --- | --- | --- | --- | --- | --- | --- | --- | --- |
| JB001 | Candidatus_Nanogingivalaceae_FGB1_strain_JCVI_27_bin.3 | 67 | 704,215 | 1 | 662,051 | 662,051 | * |  |
| JB002 | Candidatus_Saccharimonas_sp._strain_JCVI_32_bin.49 | 14 | 620,057 | 1 | 639,737 | 639,737 | * |  |
| JB003 | Candidatus_Nanogingivalaceae_FGB1_strain_JCVI_28_bin.11 | 34 | 719,702 | 1 | 663,171 | 663,171 | * |  |
| TM7c-JB | Candidatus_Nanosynbacter_TM7c_strain_JCVI_32_bin.19 | 7 | 793,808 | 1 | 793,363 | 793,363 |  | * |
| none | Candidatus_Nanosynbacter_sp._TM7_MAG_III_A_2_strain_JCVI_32_bin.12 | 76 | 696,341 | 8 | 837,467 | 808,188 |  | * |
| none | Candidatus_Nanosynbacter_GGB2_strain_JCVI_32_bin.57 | 32 | 1,040,784 | 6 | 1,054,499 | 762,750 |  | * |
| G6_32_bin_33_unicycler | Candidatus_Nanogingivalaceae_FGB1_strain_JCVI_32_bin.33 | 97 | 521,278 | 31 | 594,688 | 77,761 |  |  |
| none | Candidatus_Nanosynbacteraceae_FGB1_strain_JCVI_32_bin.22 | 68 | 636,728 | 35 | 913,508 | 182,700 |  |  |
| none | Candidatus_Nanosynbacteraceae_FGB2_strain_JCVI_32_bin.44 | 31 | 725,781 | 15 | 819,428 | 300,554 |  |  |
| G3_32_bin_36_unicycler | Candidatus_Nanosyncoccus_FGB2_strain_JCVI_32_bin.36 | 32 | 667,180 | 4 | 688,219 | 265,262 |  | * |

Phylogenetic analysis using concatenated protein sequences was performed using Anvi’o (12), and included the 8 improved/completed genomes from this study, all 26 complete Saccharibacteria genomes available on NCBI (as of 1 April 2021), and 90 Saccharibacteria draft genomes from 5 recent studies (Table S1). JB001, JB002, and JB003 were indeed members of Saccharibacteria clade G6 (Figure 1A, Figure S1), and represent the only human-associated, complete Saccharibacteria genomes outside of clade G1. Notably, G6 had the smallest genomes and the lowest GC-content of all Saccharibacteria (Figure 1A). Percent average nucleotide identity (ANI) between the G6 genomes was calculated using Anvi’o and suggested that there are at least 3-5 distinct species within the clade (Figure 1B; a cutoff of 95% ANI is frequently used to estimate the species level (13, 14)). JB001, JB003, JCVI_1_bin.12, and G6_32_bin_33_unicycler appear to be the same species, with an ANI of ≥95%, despite their source from different human subjects and independent genome assembly (Figure 1B). JB002 and T-C-M-Bin-00022 were over 98% ANI, likely representing the same distinct species, while CMJM-G6-HOT-870 and T-C-M-Bin-00011 were ∼98% ANI and formed what is likely an additional G6 species (Figure 1B). CLC Genomics Workbench was used to perform whole genome alignment for JB001, JB002, JB003, and the G1 reference strain, TM7x (Figure 1C). While JB001 and JB003 were completely syntenic, and there were moderate differences between JB001/JB003 and JB002, TM7x and the G6 Saccharibacteria have undergone many genomic re-arrangements and instances of gene gain/loss since their last common ancestor (Figure 1C).

**Figure 1:**
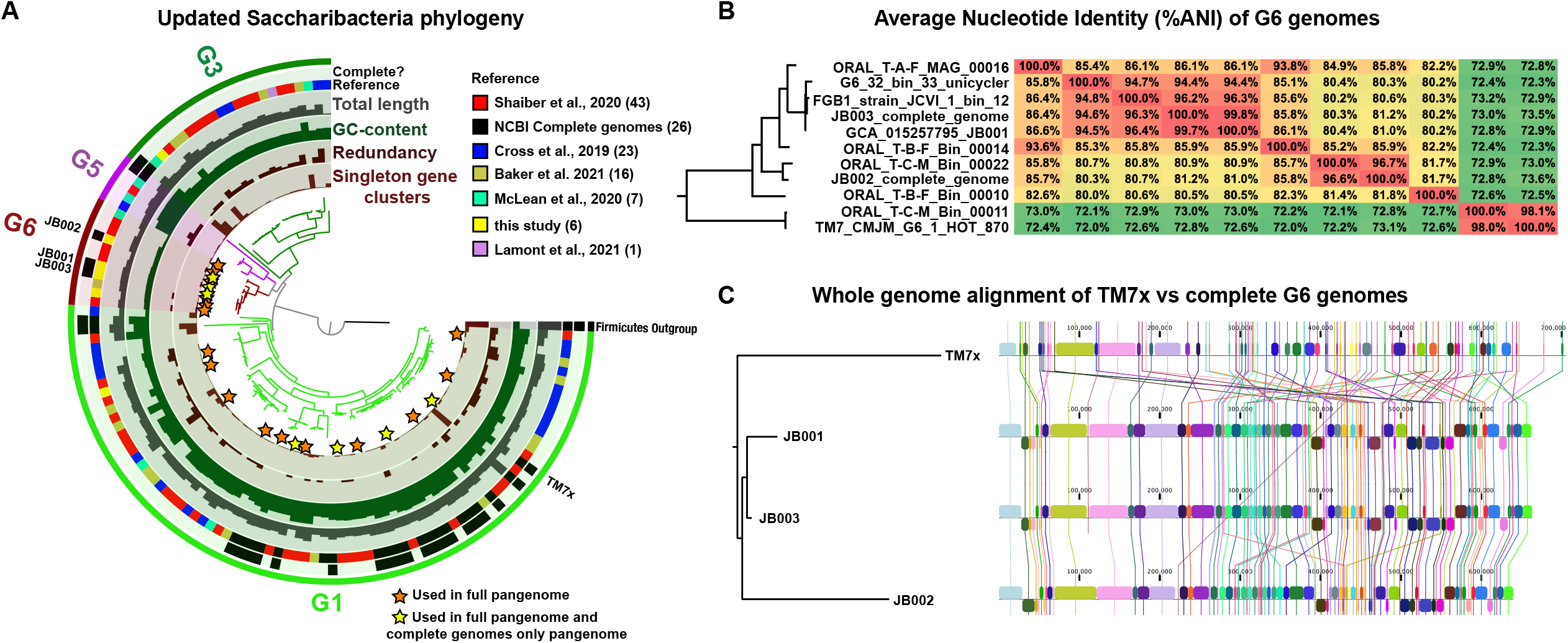
**JB001, JB002, and JB003 are clade G6 Saccharibacteria representing two distinct species. (A) Phylogenetic tree of Saccharibacteria annotated with genome data**. Phylogenetic analysis of the 123 Saccharibacteria genomes listed in Table S1. Firmicutes was used as an outgroup. The bars in the innermost layer represent the number of singleton gene clusters (i.e. genes appearing in only that one genome) in each genome. The bars in the second layer represent the redundancy (likely contamination) within each genome. The bars in the third layer represent the %GC content of each genome. The bars in the fourth layer represent the total length in bp of each genome. The fifth layer displays the source/reference for each genome. The sixth layer displays the genomes that are complete. The outermost layer, and the color of the branches of the tree, illustrate which Saccharibacteria clade each genome is part of. Orange stars indicate genomes that were used in the full pangenome analysis (Figure S2, Table S4). Yellow stars indicate genomes that were used in the pangenome analysis of compete genomes only (Figure 2, Table S3) as well as the full pangenome analysis (Figure S2, Table S4). A larger version of this figure, with the name of each genome labeled, is available in Figure S1. Note that CP025011_1_Candidatus_Saccharibacteria_bacterium_YM_S32_TM7_50_20_chromosome_c omplete_genome and c_000000000001 (GCA_003516025.1_ASM351602v1_genomic.fa), the only two complete genomes in clades G3 and G5, are from environmental, not oral, samples. The raw data in the annotations of the tree is available in Table S1. **(B) Average nucleotide identity (%ANI) of G6 genomes**. Heatmap of all-vs-all comparison of %ANI of all 11 G6 genomes. The tree on the right is a scaled up version of the G6 portion of the phylogenetic tree in panel A. Full percentage identity, which takes alignment length into account, is available in Table S2. **(C) Whole genome alignment of TM7x vs complete G6 genomes**. Whole genome alignment diagram produced by CLC Genomics Workbench. The tree on the right is based on the whole genome alignment itself.

To examine functional and metabolic differences between the G6 clade and the more well-understood G1 clade, pangenome analysis was performed using Anvi’o (15) on the 3 complete G6 genomes and 4 diverse G1 complete genomes (Figure 2, Table S3). This identified 223 “pan-Saccharibacteria Core Genes” appearing in all genomes, as well as all 94 “G1 Core Genes”, and 244 “G6 Core Genes” (Figure 2A). While 97% of the pan-Saccharibacteria Core Genes and 89% of the G1 Core Genes had known COG functions and pathways, only 50% of the G6 Core Genes had known COG functions and pathways (Figure 2A), highlighting the enigmatic nature of this clade. The likely reason for the lower number of G1 core genes is the larger amount of known diversity within the G1 clade and the genomes analyzed here (8, 9), leading to less conservation across the G1 pangenome. A larger pangenome analysis, examining all 11 G6 genomes and 14 diverse G1 genomes is available in Figure S2 and Table S4. This generated similar results, but note that this analysis contains incomplete draft genomes which are incomplete and/or may contain contamination. A complete metabolic network illustrating the known KEGG pathways identified in the three sets of core genes identified in Figure 2A is shown in Figure 2B. Both G1 and G6 genomes encode partial cell wall metabolism, glycolysis (missing phosphofructokinase), and arginine biosynthesis pathways, and do not encode fatty acid metabolism, a TCA cycle, or amino acid metabolism (other than arginine) (Figure 2B). Notable pathways present in G6 genomes but absent in G1 include: maltase glucoamylase (to metabolize starch), fructose bisphosphate aldolase (a glycolytic step), adenylate cyclase, lactate dehydrogenase, partial lipoarabinomannan (LAM) biosynthesis, and partial glycerolipid metabolism. Conversely, G1 genomes encode the non-oxidative phase of the pentose phosphate pathway, an F1F0 ATPase, alpha galactosidase, and several steps in nucleotide metabolism, which were not present in the G6 genomes (Figure 2B). Between JB001 and JB002, most differences were genes with unknown functions, therefore the differences in the KEGG pathways encoded were minor (Figure S3). The G6 genomes examined did not contain predicted elements of a CRISPR system. Although it is not known how Saccharibacteria obtain needed metabolites from the host, a type IV pilus-like system is generally well-conserved across the group, has been proposed as a candidate mechanism (8, 9), and was present in the G6 genomes here. The species-level clade that included JB001 and JB003 encoded a ∼10,000bp putative prophage element, which was flanked by homologs to the PinE invertase and contained a T4SS VirD4 homolog and 4 hypothetical proteins, all with ∼95% homology to a similar region in *Streptococcus salivarius*.

**Figure 2:**
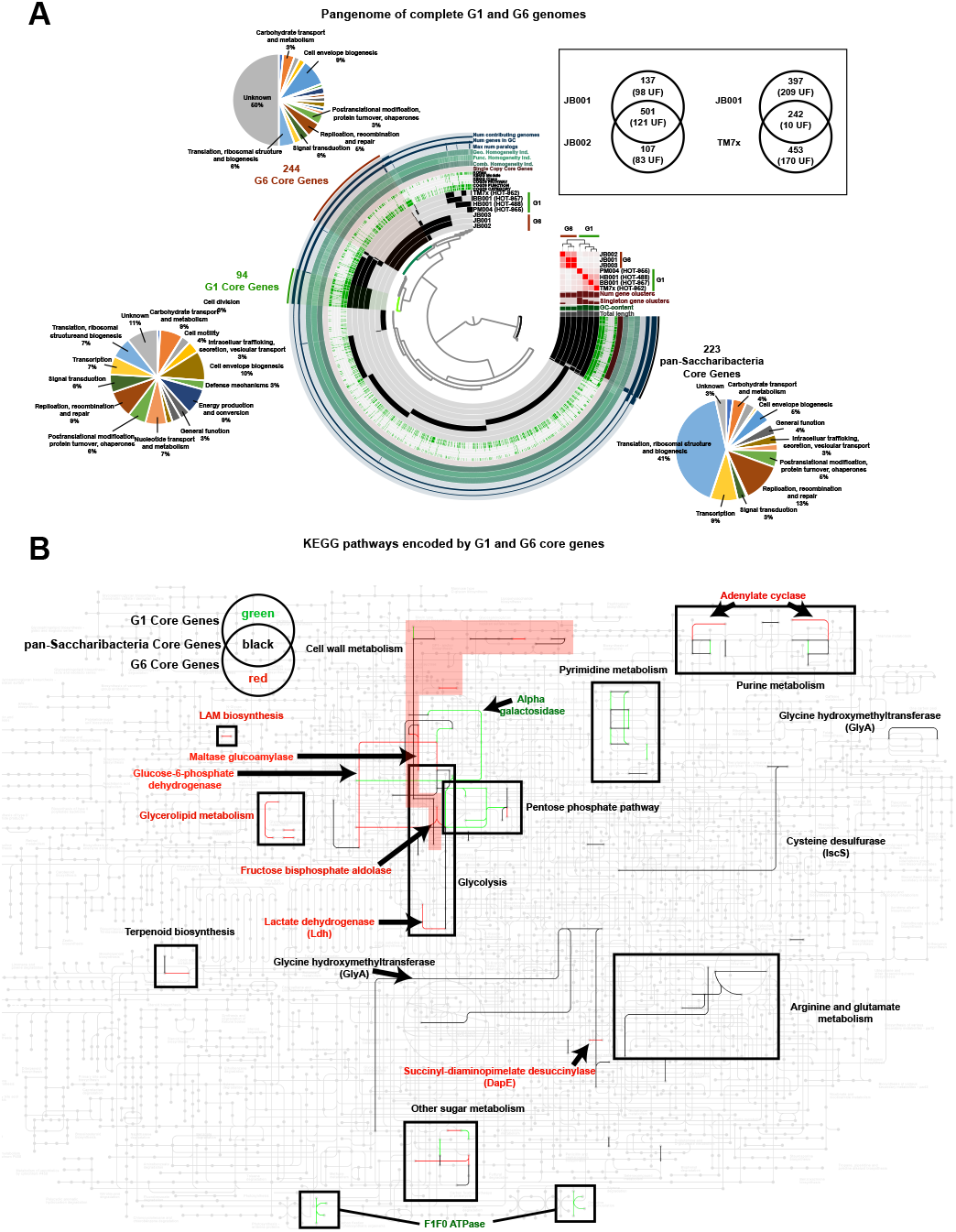
**Pangenome analysis of complete genomes in Saccharibacteria clade G1 vs. clade G6 identifies core genes with encoding distinct functional pathways. (A) The pangenome of complete G1 and G6 genomes**. The dendrogram in the center organizes the 2,279 gene clusters identified across in the genomes represented by the innermost 7 layers: TM7x, BB001, HB001, PM004, JB003, JB001, and JB002. The data points within these 7 layers indicate the presence of a gene cluster in a given genome. From inside to outside, the next 6 layers indicate known vs unknown COG category, COG function, COG pathway, KEGG class, KEGG module, and KOfam. The next layer indicates single-copy pan-Saccharibacteria core genes. The next 6 layers indicate the combined homogeneity index, functional homogeneity index, geometric homogeneity index, max number of paralogs, number of genes in the gene cluster, and the number of contributing genomes. The outermost layer highlights gene clusters that correspond to the pan-Saccharibacteria Core Genes (found in all 7 genomes), the G1 Core Genes (found in all G1 genomes and no G6 genomes), and the G6 Core Genes (found in all G6, but no G1 genomes). The pie chart adjacent to each group of core genes indicates the breakdown of COG categories of the gene clusters in the group. The 7 genome layers are ordered based on the tree of the %ANI comparison, which is displayed with the red and white heatmap. The layers underneath the %ANI heatmap, from top to bottom, indicate: the number of gene clusters, the number of singleton gene clusters, the GC-content, and the total length of each genome. The Venn diagrams in the inset show the number of overlapping and non-overlapping genes between JB001 and JB002, and JB001 and TM7x. The number in parenthesis is the number of genes with unknown functions (UF). **(B). KEGG pathways encoded by G1 and G6 core genes**. KEGG **m**etabolic map overlaid with the pathways encoded by the pan-Saccharibacteria core genes (black), G1 Core Genes (green), and G6 Core Genes (red), as indicated by the Venn diagram key. Enzymes of interest are labeled with text and arrows. Pathways are indicated by labeled boxes, the cell wall metabolism pathways is labeled with the red background to distinguish it due to the odd shape and overlap with the glycolysis pathway space.

Taken together, these analyses indicate that Saccharibacteria clade G6 is highly divergent from clade G1, and may have a different lifestyle, host, and host-dependencies. This is in line with the recent hypothesis that G6 reside on the tongue (G6 are referred to as ‘T2’ in reference 9) and have a long history of association with animal hosts, while G1 reside in dental plaque and were a much more recent acquisition from the environment (8, 9). Interestingly, the species-level clade containing JB002 (the most reduced Saccharibacteria genome, with only 615 genes) was the only Saccharibacteria group that resided both on the tongue and in dental plaque (9). Although all cultured isolates of Saccharibacteria were epibionts of *Actinomyces* spp., they were all G1 strains. Residing in a different environment, G6 may have distinct host species, possibly *Streptococcus*, given the acquired homologous sequence. It is likely that G6 fallen into the ‘unknown’ taxonomic bucket in the majority of past microbiome studies, thus the role of G6 in human health remains to be elucidated. The high percentage of genes with unknown functions further adds to the obscurity of this clade. Overall, this article highlights an urgent need for study of Saccharibacteria, since almost nothing is known about the lifestyle, host, or ecological impact of Saccharibacteria clade G6, and even less still is understood about clades G2, G3, G4, and G5.

## Acknowledgements

I thank Karrie Goglin-Almeida, Jelena Jablanovic, and Kara Riggsbee for performing the library preparation and sequencing, and Jeffrey S. McLean for helpful discussions. This research was supported by NIH/NIDCR K99-DE029228.

## Data availability

The complete genome sequences of JB001, JB002, and JB003 have been deposited in GenBank under the accession numbers: <u>CP072208</u>, <u>CP076101</u>, and <u>CP076102</u>. The BioProject accession for this project is <u>PRJNA624185</u>. The short reads used to generate the assemblies are available in the SRA database with the accession numbers <u>SRX4318838, SRX4318837</u>, and <u>SRX4318835</u>. The long reads used to generate the assemblies are available in the SRA dataset with the accession numbers <u>SRX10387815</u>, <u>SRX11020560</u> and <u>SRX11020561</u>.

