## Supplemental Material for "Complete genomes of clade G6 Saccharibacteria suggest a divergent ecological niche and lifestyle"

#### Supplemental methods

**Nanopore sequencing.** A recent study (1) provided a number of Saccharibacteria draft genomes that were of particular interest due to their high quality compared to the other genomes from those clades available in public databases (high N50, low number of contigs, low contamintion) (Table 1). Nanopore sequencing was used to close the genome of JB001, a member of Saccharibacteria clade G6. The methods used to obtain the JB001 genome are described in the JB001 *Microbiology Resource Announcement* (in press). Here, similar methods, described below, were used to improve the genome assembly of 9 additional Saccharibacteria genomes. Eight of these genomes: Candidatus_Saccharimonas_sp._strain_JCVI_32_bin.49, Candidatus_Nanosynbacter_TM7c_strain_JCVI_32_bin.19, Candidatus_Nanosynbacter_sp._TM7_MAG_III_A_2_strain_JCVI_32_bin.12, Candidatus_Nanosynbacter_GGB2_strain_JCVI_32_bin.57, Candidatus_Nanogingivalaceae_FGB1_strain_JCVI_32_bin.33, Candidatus_Nanosynbacteraceae_FGB1_strain_JCVI_32_bin.22, Candidatus_Nanosynbacteraceae_FGB2_strain_JCVI_32_bin.44, and Candidatus_Nanosyncoccus_FGB2_strain_JCVI_32_bin.36 were originally assembled from an ultra-deep (~300M reads) Illumina read library representing the oral microbiome of a single human subject with healthy teeth, SC33 (NCBI accession number SRR7448307) (1). Meanwhile, the G6 genome Candidatus_Nanogingivalaceae_FGB1_strain_JCVI_28_bin.11 was originally assembled from an Illumina read library representing the oral microbiome of a different human subject with healthy teeth, SC24 (NCBI accession number SRR7449305) (1). In an attempt to close these 9 genomes, Oxford Nanopore sequencing was performed on the same two saliva samples used to generate the original short-read assemblies (1). High molecular weight genomic DNA (HMW gDNA) was extracted using a previously described phenol-chloroform based method

(2). The resulting gDNA was examined for purity, size, and concentration using a TapeStation (Agilent Technologies) and a Qubit fluorometer (Thermo Fisher Scientific). DNA was not sheared or size-selected. A long-read library was prepared using a Ligation Sequencing Kit (Oxford Nanopore Technologies) and sequenced on a GridION using an R9.4.1 Flow Cell (Oxford Nanopore Technologies). Basecalling, adapter trimming, and quality control were performed using Guppy v4.8.11/MinKNOWv20.06.9.

**Genome reassembly.** Long reads mapping to each of the 9 genomes of interest from Baker et. al (1) were extracted using minimap2 v2.17-r941 and samtools v1.7. The original draft genomes had been assembled using the MetaWRAP pipeline (3), and therefore had been reassembled using only reads that mapped to the first assembly, and the reassembled genome had been used going forward in the cases where reassembly improved completeness and reduced contamination (i.e. the Reassemble_bins module of MetaWRAP) (3). For each genome, the "permissive" set of reads mapping to each genome identified by MetaWRAP was used, along with the extracted mapped long reads generated by the nanopore sequencing, to create a new assembly using Unicycler v0.4.8 (4). The resulting Unicycler assembly for each genome, and additional steps to attempt to improve and close each assembly are discussed here, genome by genome:

**JB002 (G6)**. Assembly with Unicycler resulted in 3 contigs, one circular contig of 638,658 bp, and two linear contigs of 1,294 bp and 1,072 bp. Upon further inspection using Anvi'o v7-dev (5), the two short contigs were removed based on disparate coverage, tetranucleotide frequency, and blast hits to other organisms. To examine the genome for problems, CLC genomics Workbench 21 was used to map the long and short read sets to the circular contig and visualize coverage. There were 6 low coverage regions (< 5X coverage) totally 7,659 bp. There may be reads in the libraries that map to the updated assembly, but did not map to the original assembly, and might therefore be used to improve the current assembly. Reads from the complete, original short read library were mapped to the updated assembly using bwa v0.7.17-r1188 and reads from the long-read library were mapped with minimap2. Fastq files of mapped reads were extracted with samtools, and QC of the mapped short reads was performed with fastp v0.20.1. These updated read sets were assembled using Unicycler. This time, 72 contigs were produced, however the largest contig was one that was circular and 639,737 bp. This time, CLC analysis of read mappings to the circular contig revealed only 5 areas of low coverage, totally only 327 bp, which were all in the 112,000-119,000 bp region. Circulator v.1.5.5 (6) was used to rotate the genome start to the *dnaA* gene. The completed genome was annotated using the NCBI Prokaryotic Genome Annotation Pipeline (PGAP) v5.1.

**JB003 (G6).** The Unicycler assembly produced 7 linear contigs, however 1 was 691,752 bp and the rest were under 10,000 bp in length. Examination using Anvi'o revealed that the short contigs had disparate coverage and tetranucleotide frequency, and had blast hits to other organisms, therefore they were removed. A remapping of the read libraries to the updated assembly was performed as described above for JB002. The 2$^{nd}$ iteration of Unicycler, using the updated read sets, resulted in 6 contigs, with the largest being 663,171 bp and circular. Here, there was a 27,975 bp circular contig and 3 short contigs of ~1,500 bp. Again, the short contigs were identified as assembly artifacts by Anvi'o and removed. There were no problematic regions of low coverage identified by CLC Genomics Workbench on the closed genome. Circulator v.1.5.5 (6) was used to rotate the genome start to the *dnaA* gene. The completed genome was annotated using the NCBI Prokaryotic Genome Annotation Pipeline (PGAP) v5.1.

**G6_32_bin_33_unicycler (G6).** Unicycler assembly resulted in 31 contigs, all were linear and the longest was 594,688 bp. Since this genome was not close to complete, no further attempts were made to close this genome.

**G3_32_bin_36_unicycler (G3).** Assembly with Unicycler resulted in 4 contigs, all linear, with the largest being 265,262 bp and others being 235,441 bp, 173,991 bp, and 13,325 bp.. Remapping to the updated assembly and reassembly with the mapped reads did not improve this assembly. With 4 large contigs, and an assembly size that would make sense for a G3 Saccharibacteria (although no complete G3 genomes have been reported) this assembly is "near complete." Notably, this is currently by far the most contiguous and highest quality G3 genome obtained to date.

**TM7c-JB (G1).** Unicycler assembly resulted in 11 contigs. The largest was 793,363 bp and linear. Based upon different GC content, tetranucleotide frequency, and blast hits to other organisms (performed using Anvi'o), the 10 smaller contigs were identified as contamination and removed. A remapping approach, similar to above for JB002 and JB003, was attempted, but did not circularize the contig or further improve the assembly. Since the initial longest contig is the expected size for a G1 TM7 and appears to only have one gap, this genome is "near complete."

**Candidatus_Nanosynbacter_sp._TM7_MAG_III_A_2_strain_JCVI_32_bin.12 (G1).** Unicycler assembly resulted in 8 contigs, with the largest being linear and 808,188 bp. The 7 short contigs were identified as contamination using Anvi'o and removed. Remapping and reassembly did not circularize or improve the assembly, which remains "near complete" as 808,188 bp is an expected size for a G1 Saccharibacteria.

**Candidatus_Nanosynbacter_GGB2_strain_JCVI_32_bin.57 (G1).** Unicycler assembly resulted in 6 contigs, with the largest being 762,750 bp. Although the largest contig is "near complete" with an expected size of a G1 Saccharibacteria, examination with Anvi'o could not discern if the other contigs were indeed contamination. Furthermore, remapping and reassembly did not improve the genome. Together, the 6 contigs totaled 1,054,499 bp, which is probably too large for an oral G1 Saccharibacteria and therefore the full assembly here likely represents a composite genome of multiple G1 strains.

**Candidatus_Nanosynbacteraceae_FGB1_strain_JCVI_32_bin.22 (G1).** Unicycler assembly resulted in 35 contigs, with the largest being 182,700 bp. Since this genome was not close to complete, no further attempts were made to close this genome.

**Candidatus_Nanosynbacteraceae_FGB2_strain_JCVI_32_bin.44 (G1).** Unicycler assembly resulted in 15 contigs, with the largest being 300,554 bp. Since this genome was not close to complete, no further attempts were made to close this genome.

**Phylogenomics.** Phylogenomics was performed on the 123 Saccharibacteria genomes listed in Table S1 using the Anvi'o Phylogenomics Snakemake (7) Workflow using the default settings with the exception that alignment was performed using muscle instead of FAMSA and the number of threads available was increased to 50. The phylogenetic trees in Figure 1A, Figure 1B and Figure S1 were viewed and annotated using the anvi-interactive script.

**Genome similarity (average nucleotide identity [ANI]).** Genome similarity was determined for the 11 G6 genomes using Anvi'o. The resulting text files with the data for %ANI (Figure 1B) and full percentage identity (which takes alignment length into account) (Table S2) were reordered based upon the phylogenomic tree and turned into a heatmap using Microsoft Excel v16.16.23.

**Whole genome alignment.** Whole genome alignment was performed and visualized using CLC Genomics Workbench 21 using the default settings.

**Pangenome analysis.** Pangenome analysis was performed on either the 3 complete G6 genomes and 4 select, complete G1 genomes ("complete genomes only pangenome", genomes analyzed indicated by yellow stars in Figure 1A) (Figure 2A) OR all 11 G6 genomes and 14 diverse, select G1 genomes ("full pangenome", genomes analyzed indicated by yellow and orange stars in Figure 1A) (Figure S2) using the Anvi'o Pangenomics Snakemake Workflow. Default settings were used except that alignment was performed using muscle instead of FAMSA and the number of threads available was increased to 50. The pangenomes were displayed using the Anvi'o anvi-display-pan script and the "pan-Saccharibacteria Core Genes" (present in all genomes), "G1 Core Genes" (present in all G1 but no G6), and "G6 Core Genes" (present in all G6 but no G1) bins were manually selected. Data from the pangenomes was exported into tabular text format (Tables S3 and S4) using the anvi-summarize command. The pangenome information was used to create the Venn diagrams in the Figure 2A inset, and Microsoft Excel was used to create the COG pathway pie charts for each of the 3 core gene bins. The KEGG KO identifiers from the pangenome data were used as input for the KEGG Mapper Search&Color pathway tool (https://www.genome.jp/kegg/mapper.html), which was used to generate the metabolic networks used in Figure 2B and Figure S3.

**Supplemental figure legends.**

**Figure S1: JB001, JB002, and JB003 are clade G6 Saccharibacteria representing two distinct species. (A) Phylogenetic tree of Saccharibacteria annotated with genome data (the same as Figure 1 in the main text except all leaves of the phylogenetic tree are labeled).** Phylogenetic analysis of the 123 Saccharibacteria genomes listed in Table S1. Firmicutes was used as an outgroup. The bars in the innermost layer represent the number of singleton gene clusters (i.e. genes appearing in only that one genome) in each genome. The bars in the second layer represent the redundancy (likely contamination) within each genome. The bars in the third layer represent the %GC content of each genome. The bars in the fourth layer represent the total length in bp of each genome. The fifth layer displays the source/reference for each genome. The sixth layer displays the genomes that are complete. The outermost layer, and the color of the branches of the tree, illustrate which Saccharibacteria clade each genome is part of. Orange stars indicate genomes that were used in the full pangenome analysis (Figure S2, Table S4). Yellow stars indicate genomes that were used in the pangenome analysis of compete genomes only (Figure 2, Table S3) as well as the full pangenome analysis (Figure S2, Table S4). A larger version of this figure, with the name of each genome labeled, is available in Figure S1. Note that CP025011_1_Candidatus_Saccharibacteria_bacterium_YM_S32_TM7_50_20_chromosome_complete_genome and c_000000000001 (GCA_003516025.1_ASM351602v1_genomic.fa), the only two complete genomes in clades G3 and G5, are from environmental, not oral, samples. The raw data in the annotations of the tree is available in Table S1. **(B) Average nucleotide identity (%ANI) of G6 genomes.** Heatmap of all-vs-all comparison of %ANI of all 11 G6 genomes. The tree on the right is a scaled up version of the G6 portion of the phylogenetic tree in panel A. Full percentage identity, which takes alignment length into account, is available in Table S2. **(C) Whole genome alignment of TM7x vs complete G6 genomes.** Whole genome alignment

**Figure S1**

#### Saccharibacteria Phylogeny

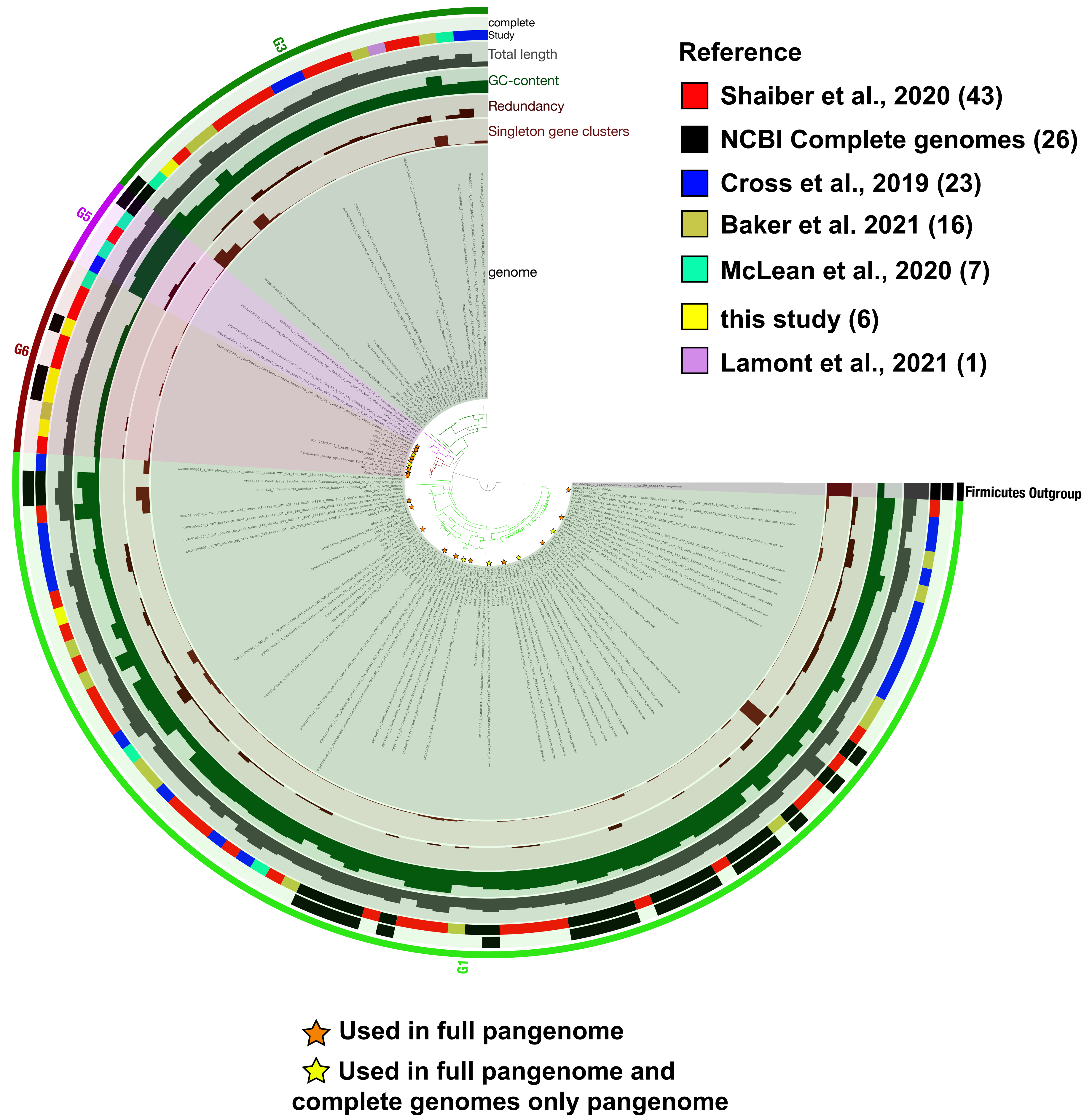

diagram produced by CLC Genomics Workbench. The tree on the right is based on the whole genome alignment itself.

**Figure 2: Pangenome analysis of all 11 genomes in Saccharibacteria clade G6 vs. 14 diverse G1 genomes identifies core genes with encoding distinct functional pathways.** The dendrogram in the center organizes the 4,452 gene clusters identified across in the genomes represented by the innermost 25 layers. The data points within these 25 layers indicate the presence of a gene cluster in a given genome. From inside to outside, the next 6 layers indicate known vs unknown COG category, COG function, COG pathway, KEGG class, KEGG module, and KOfam. The next layer indicates single-copy pan-Saccharibacteria core genes. The next 6 layers indicate the combined homogeneity index, functional homogeneity index, geometric homogeneity index, max number of paralogs, number of genes in the gene cluster, and the number of contributing genomes. The outermost layer highlights gene clusters that correspond to the pan-Saccharibacteria Core Genes (found in all genomes), the G1 Core Genes (found in all G1 genomes and no G6 genomes), and the G6 Core Genes (found in all G6, but no G1 genomes). The 25 genome layers are ordered based on the tree of the %ANI comparison, which is displayed with the red and white heatmap. The layers underneath the %ANI heatmap, from top to bottom, indicate: the Saccharibacteria clade, the number of gene clusters, the number of singleton gene clusters, the number of gene clusters per kbp, the redundancy (i.e. probable contamination), the completion, the GC-content, and the total length of each genome. **(B). KEGG pathways encoded by G1 and G6 core genes.** KEGG metabolic map overlaid with the pathways encoded by the pan-Saccharibacteria core genes (black), G1 Core Genes (green), and G6 Core Genes (red), as indicated by the Venn diagram key. Enzymes of interest are labeled with text and arrows. Pathways are indicated by labeled boxes.

### Figure S2

## A

##### G6vsG1 full pangenome

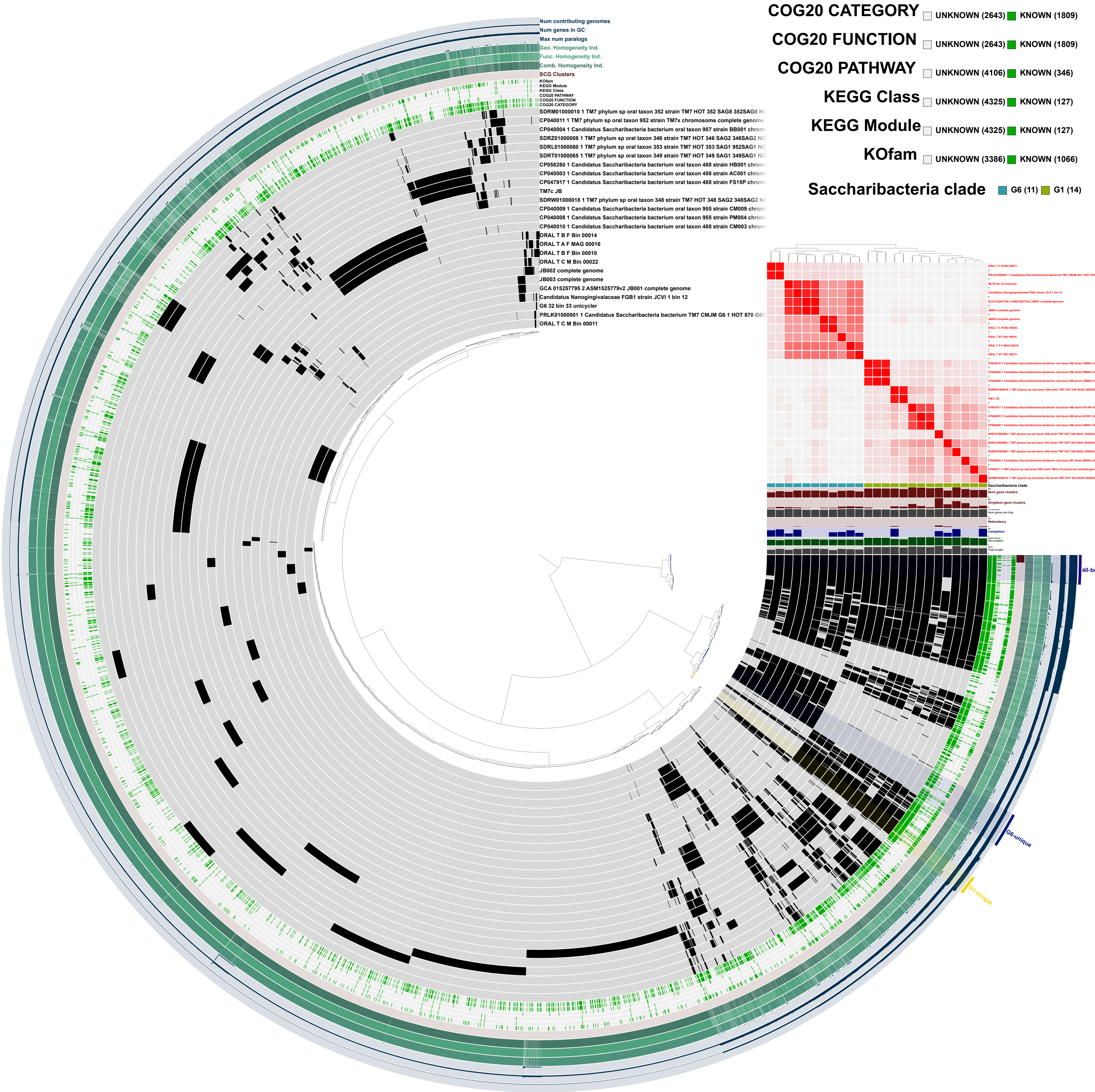


## B

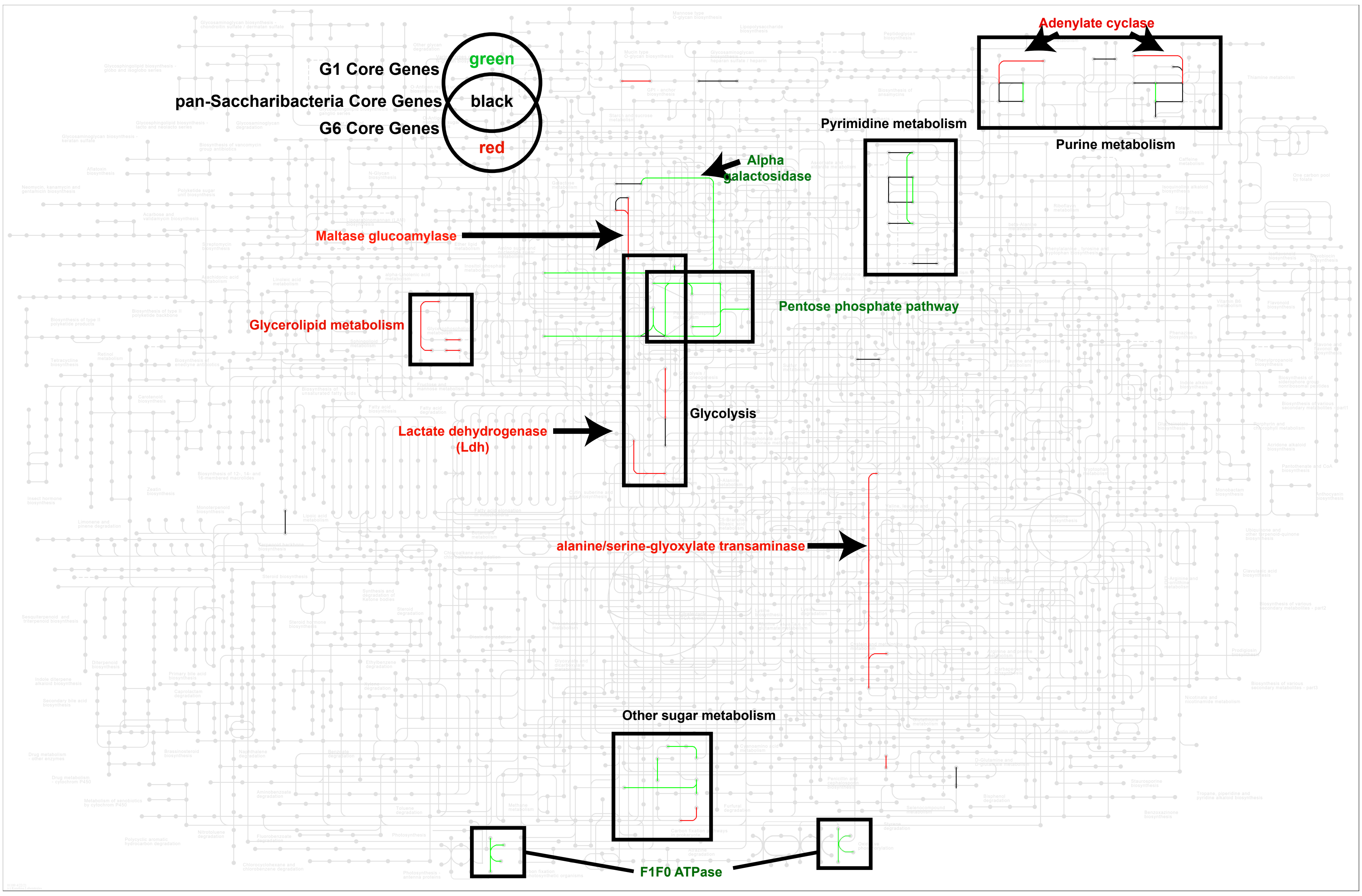

**Figure S3: Differences in KEGG pathways encoded by JB001 and JB002.** KEGG metabolic map overlaid with the pathways encoded both JB001 and JB002 (black), JB001 only (red), and JB002 only (green), as indicated by the venn diagram key. Enzymes of interest are labeled with text and arrows. Pathways are indicated by labeled boxes.

#### Figure S3

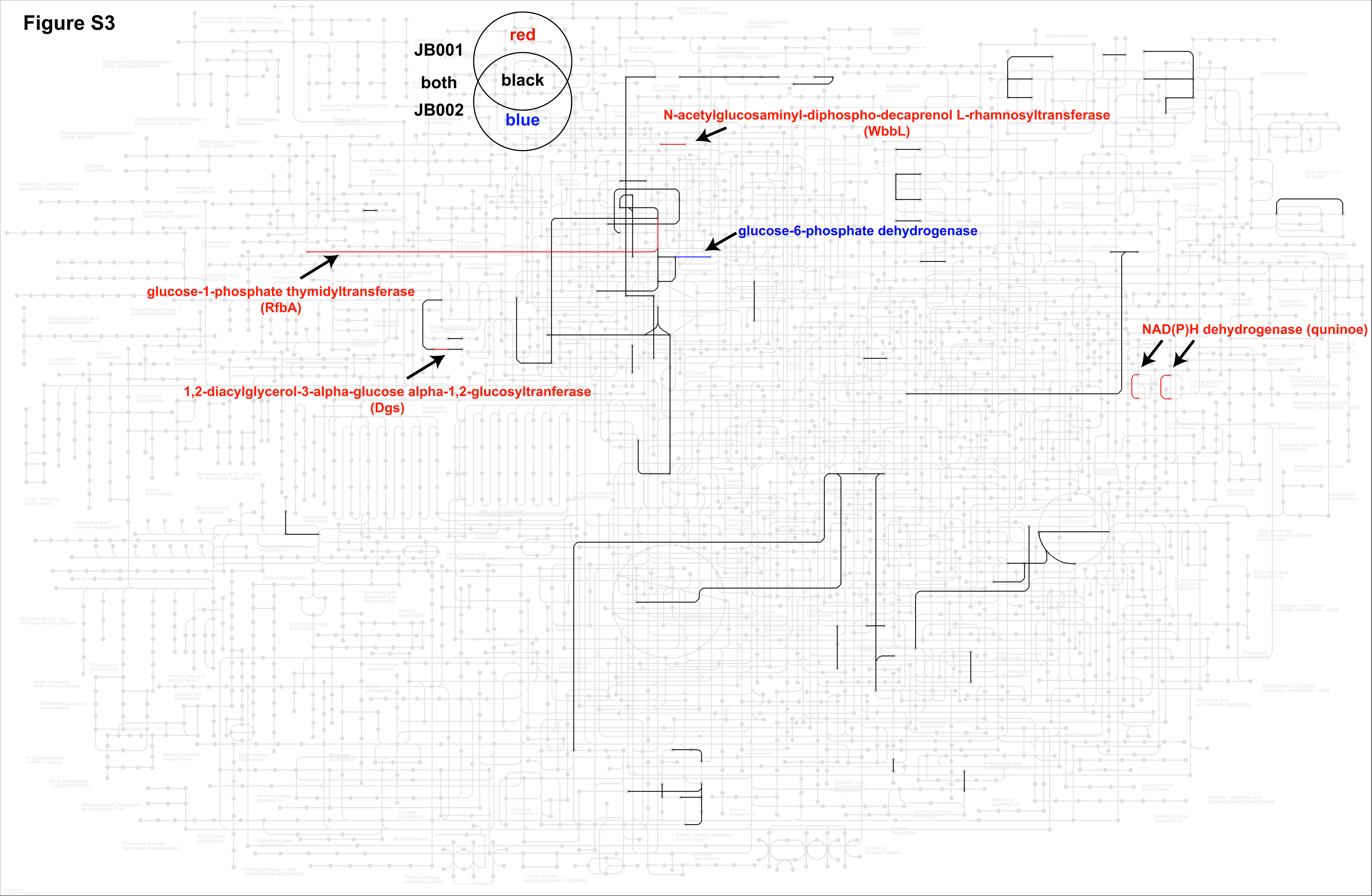
JB001
both
JB002
red
black
blue
N-acetylglucosaminyl-diphospho-decaprenol L-rhamnosyltransferase (WbbL)
glucose-6-phosphate dehydrogenase
glucose-1-phosphate thymidyltransferase (RfbA)
1,2-diacylglycerol-3-alpha-glucose alpha-1,2-glucosyltranferase (Dgs)
NAD(P)H dehydrogenase (quninoe)
